# Differential Functions of Splicing Factors in Breast-Cancer Initiation and Metastasis

**DOI:** 10.1101/634154

**Authors:** Shipra Das, Martin Akerman, SungHee Park, Mattia Brugioli, Adam Geier, Anil K. Kesarwani, Martin Fan, Nathan Leclair, Laura Urbanski, Kuan-Ting Lin, Chenle Hu, Xingan Hua, Joshy George, Senthil K. Muthuswamy, Adrian R. Krainer, Olga Anczuków

**Affiliations:** Cold Spring Harbor Laboratory, Cold Spring Harbor, NY, USA; Envisagenics, Inc, New York, NY, USA; The Jackson Laboratory for Genomic Medicine, Farmington, CT, USA; Graduate Program in Genetics and Development, UConn Health, Farmington, CT, USA; Beth Israel Deaconess Medical Center, Boston MA 02215, USA

**Author notes:** Equal contributions (SungHee Park, Mattia Brugioli).

## Abstract

Misregulation of alternative splicing is a hallmark of human tumors; yet to what extent and how it contributes to malignancy are only beginning to be unraveled. Here, we define which members of the splicing factor SR and SR-like families contribute to breast cancer, and uncover differences and redundancies in their targets and biological functions. We first identify splicing factors frequently altered in human breast tumors, and then assay their oncogenic functions using breast organoid models. Importantly we demonstrate that not all splicing factors affect mammary tumorigenesis. Specifically, upregulation of either SRSF4, SRSF6 or TRA2β promotes cell transformation and invasion. By characterizing the targets of theses oncogenic factors, we identify a shared set of spliced genes associated with well-established cancer hallmarks. Finally, we demonstrate that the splicing factor TRA2β is regulated by the MYC oncogene, plays a role in metastasis maintenance *in vivo*, and its levels correlate with breast-cancer-patient survival.

## INTRODUCTION

Cancers arise as a consequence of the dysregulation of cellular homeostasis and its multiple control mechanisms. Alternative RNA splicing is a key step in gene-expression regulation and contributes to transcriptional diversity and plasticity by selecting which transcript isoforms are produced in a specific cell at a given time. Defects in alternative splicing are frequently found in human tumors, and RNA-splicing regulators have recently emerged as a new class of oncoproteins or tumor suppressors (Dvinge et al., 2016). Alterations in RNA splicing can lead to malignancy by affecting the expression of oncogene and tumor-suppressor isoforms (Biamonti et al., 2014). Tumor-associated aberrant alternative-splicing profiles result either from mutations in splicing-regulatory elements of specific cancer genes or from changes in the splicing machinery (Urbanski et al., 2018). Recurrent somatic mutations in selected components of the splicing machinery frequently occur in myeloid tumors, suggesting that splicing-factor (SF) alterations are a hallmark of cancer (Yoshida and Ogawa, 2014). In solid tumors, SFs exhibit frequent changes in copy number and/or expression-level, but are rarely mutated (Urbanski et al., 2018).In eukaryotes, the core splicing machinery plus associated regulatory factors comprises more than 300 factors and five small nuclear RNAs, and catalyzes both constitutive and regulated alternative splicing. One major class of SFs is the serine/arginine-rich (SR) RNA-binding protein family, characterized by one or two RNA recognition motifs (RRM) and a C-terminal arginine-serine/rich (RS) domain (Manley and Krainer, 2010). SR proteins act at multiple steps of spliceosome assembly and are involved in both constitutive and alternative splicing (Black, 2003). Another group of SFs comprises the heterogeneous nuclear ribonucleoproteins (hnRNPs), which have been implicated in repressing alternative splicing events (Black, 2003). SR and hnRNP A/B proteins can exhibit antagonistic effects on the alternative splicing of particular exons. Both activator and repressor SFs can bind directly to pre-mRNA and elicit changes in alternative splicing in a concentration-dependent manner (Cáceres et al., 1994; Mayeda et al., 1992); thus, changes in SF levels are likely involved in alternative splicing deregulation in cancer even in the absence of mutations, and can affect a network of downstream targets.

Changes in alternative splicing patterns are frequently detected in human breast tumors (Eswaran et al.,2013; Venables et al., 2008; Venables et al., 2009), yet the upstream regulators controlling these tumor-associated alternatively spliced isoforms have not been extensively characterized. We previously demonstrated that overexpression of the SF SRSF1, which is frequently upregulated in human breast tumors (Karni et al., 2007), promotes transformation of mammary epithelial cells *in vitro* and *in vivo* (Anczuków et al., 2012), and acts by regulating spliced isoforms associated with cell proliferation and celldeath (Anczuków et al., 2015; Anczuków et al., 2012). SRSF1 is a prototypical member of the SR protein family, which comprises 12 members (SRSF1 to SRSF12) with structural similarities (Long and Caceres, 2009). Yet, little is known about differences and redundancies in their alternative splicing targets in human tissues, and thus in differences in their biological functions. Several lines of evidence suggest that, in addition to SRSF1, other SR proteins may play roles in breast-cancer pathogenesis: i) SR protein levels increase during murine mammary gland tumorigenesis (Stickeler et al., 1999), and, specifically, SRSF4, SRSF5, or SRSF6 are upregulated in human breast tumors or cell lines (Huang et al., 2007; Karni et al., 2007; Pind and Watson, 2003); ii) increased expression of *SRSF3* correlates with tumor grade in breast-cancer patients (Jia et al., 2010); and iii) the SR-like protein TRA2β is upregulated in human breast tumors, compared to matched normal tissue (Best et al., 2013; Watermann et al., 2006). Therefore, it is critical to characterize the differences and redundancies in SR-protein targets in cancer, as well as to assess their specificities as breast-cancer oncogenes. It remains to be determined whether dysregulation of any SF contributes to tumor initiation and/or maintenance, and what the functional consequences of SF alterations in breast cancer are. Here we describe the molecular portraits of SF alterations in human breast tumors, focusing on the SR protein family and we identify specific SF alterations that promote mammary epithelial cell transformation, using model systems relevant to the biological context in which breast tumors arise.

## RESULTS

### Comprehensive molecular portrait of SF alterations in human breast tumors

To provide a detailed understanding of SF alterations in tumors, we systematically assessed mutations, copy number changes, and expression changes in 960 human breast tumors from The Cancer Genome Atlas (TCGA) dataset (Figure 1). Based on previous studies reporting upregulation of SF levels in smaller cohorts of breast cancer patients, we focused our analysis on: i) the 12 members of the SR-protein family;ii) *TRA2β*, an SR-like protein; and iii) *HNRNPA1*, a known antagonist of the oncogenic SRSF1 factor, which was reported to be misregulated in breast tumors (Karni et al., 2007; Pelisch et al., 2012). No recurrent somatic mutations in these SFs were detected in TCGA human breast tumors (Table S1A).This is in stark contrast to the recurrent *SRSF2* mutations, centered on residue P95, detected in 20-30% of patients with myelodysplastic syndromes and 50% of those with chronic myelomonocytic leukemia (Urbanski et al., 2018). Together, these findings suggest that SF mutations are not likely to play a major role in breast cancer, in contrast to their role in myeloid tumors (Imielinski et al., 2012). Nonetheless, 57% of the 960 TCGA tumors had an alteration in at least one of the SFs analyzed, revealing that breast tumors exhibit frequent SF alterations, both in copy number and/or expression (Figure 1). Recurrent SF alterations ranged from 2 to 15% of tumors; in particular, *SRSF1, TRA2β, SRSF2*, and *SRFS6* were each altered in ≥10% of the 960 tumors, which represents a substantial proportion of breast-cancer patients, given that alterations in the known cancer drivers *ERBB2, MYC*, or *TP53* were detected in 17%, 27%, and 37% of the 960 TCGA tumors, respectively (Figure 1). Most of the SFs we examined did not show a strong correlation between copy number and expression changes in TCGA breast tumors (Table S1),suggesting additional layers of regulation, *e.g.*, at the transcriptional or post-transcriptional level. Interestingly, certain SFs were mostly upregulated, *i.e., SRSF1, SRFS2*, or *SRSF9*; others were mostly downregulated, *i.e., SRSF5* or *SRSF12;* and several SFs, including *SRSF3, SRSF4, SRFS6*, and *TRA2β* were upregulated in certain tumors and downregulated in others (Figure S1A). SFs exhibited moderate 2-3 fold changes in expression levels in the TCGA tumors (Figure S1A), in agreement with previous studies (Huang et al., 2004; Jia et al., 2010; Karni et al., 2007; Lapuk et al., 2010; Pelisch et al., 2012; Pind and Watson, 2003; Stickeler et al., 1999; Watermann et al., 2006). In addition, tumors with upregulated *SRSF1, SRSF2, SRSF3, SRSF4*, or *TRA2β* were enriched in the basal breast cancer subtype (Figure S1B), an aggressive tumor subtype. Finally, increased *TRA2β* levels were detected in 40% of triple negative breast cancer (TNBC) tumors (Figure S1C), a tumor subtype associated with poor prognosis and lack of effective therapies.

**Figure 1.**
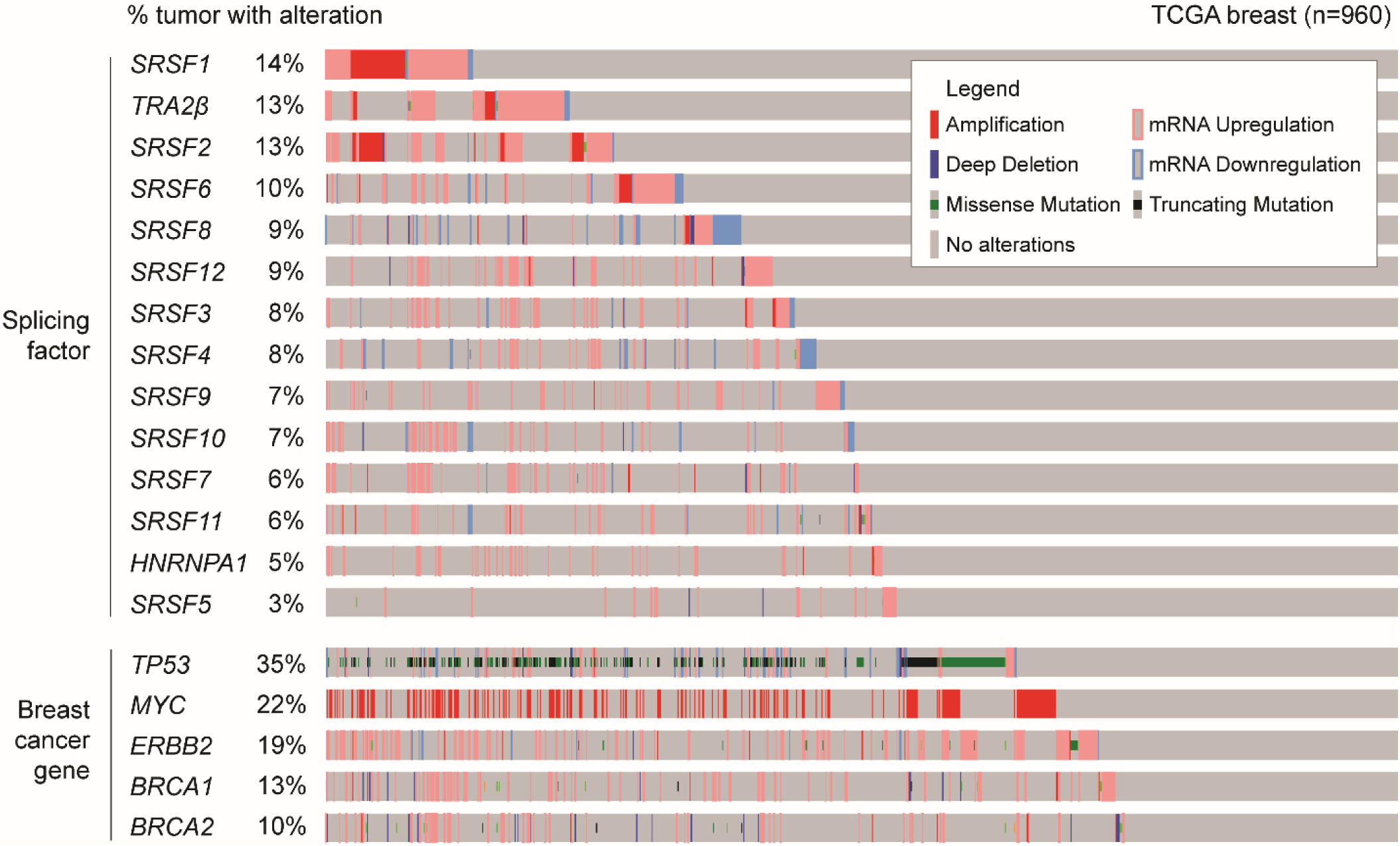
Splicing-factor alterations are detected frequently in human breast tumors. Graphical representation of SF alterations in human breast tumors from The Cancer Genome Atlas (TCGA) dataset (n=960) sorted by most frequently altered SFs. Copy number and expression changes were assessed by DNA- and RNA-seq, respectively (Cerami et al., 2012; Gao et al., 2013). Individual genes are represented as rows, and individual patients as columns. Alterations in the known breast-cancer genes *TP53, MYC*, and *ERBB2* are denoted in the lower panel.

To further determine if defects in SF levels lead to splicing-pattern alterations that could contribute to pathogenesis, we characterized the significant differentially spliced events (DSEs) in TCGA breast tumors with *SF-high vs. SF-low* levels (Figure S2 and Table S2A). We quantified alternative-splicing changes using SpliceCore^®^ (https://www.envisagenics.com/platform/), a commercial software that performs exon-centric splicing analysis by mapping RNA-sequencing (RNA-seq) data to a proprietary reference transcriptome with >5 million exon-trio models to uncover cancer-associated splicing events. We combined SpliceCore results with an in-house bioinformatics pipeline to filter and prioritize reproducible splicing changes in terms of “differential percent spliced in” scores (ΔPSI) between *SF-high vs. SF-low* tumors (See supplementary methods for details).

We focused only on DSEs detected in ≥10 tumors from both the *SF-high* and the *SF-low* groups (FigureS2B). For each of the frequently altered SF *SRSF1, SRSF2, TRA2β*, and *SRFS6* (Figure 1), we identified∼1000-1500 DSEs in *SF-high vs. SF-low* tumors (Figure S2B and Table S2B-O), and 27-47% of these DSEs were detected in ≥80 *SF-high* tumors (Figure S2C). The most frequent type of DSEs in *SRSF1-* and *TRA2β-high* tumors were cassette exon (CA) (Figure S2B), which were approximately equally divided between inclusion (ΔPSI≥10%) and skipping events (ΔPSI≤-10%) (Figure S2D). Conversely, *SRSF6-high* tumors were enriched in retained introns (RI) (Figure S2B), and the great majority of DSEs in *SRSF6-high* tumors were inclusion events (Figure S2D). Additionally, *SRSF5-* and *SRSF11-high* tumors each displayed >3,000 DSEs, with a strong increase in RI events, compared to other groups (Figure S2B). However *SRSF5-* and *SRSF11-*alterations were each detected in only 3-6% of breast tumors, and 100% of the associated DSEs were present in ≤80 *SF-high* tumors (Figure S2B,C). Finally, tumors with high levels of *SRSF3, SRSF9, SRSF12*, or *HNRNPA1* displayed ∼1000-3000 DSEs, but the majority was detected in ≤80 *SF-high* tumors (Figure S2B,C).

We conducted a pairwise comparative analysis of the 14 SF tumor groups to identify shared as well as unique DSEs associated with *SF-high* expression; shared DSEs were defined as having the same ΔPSI direction, *i.e.*, either both ΔPSI≥10% or both ΔPSI≤-10%. Our analysis revealed that 19 pairwise SF combinations shared between 20-45% of DSEs, but the remaining 72 SF pairs shared less DSEs (Table S2P). The largest overlap was detected between *SRSF3-high* and *SRSF7-high* tumors, which shared 46% of their DSEs, followed by *SRSF2-high* and *SRSF7-high* tumors sharing 31% of their DSEs, and *SRSF1-high* and *SRSF10-high* tumors sharing 31% of their DSEs (Table S2P). Among the tumors with the most frequently altered SFs, *SRSF1-high* tumors shared 21% of their DSEs with *TRA2β-high* tumors, and 12% with *SRSF2-high*; moreover, *TRA2β-high* and *SRSF2-high* tumors shared 19% of their DSEs (Table S2P). Additionally, to determine if *SF-high* tumors exhibit defects in specific gene sets, we investigated if the same genes were affected by differential splicing in distinct tumor groups; shared differentially spliced genes (DSGs) were defined as any DSE within the same gene regardless of the ΔPSI direction or the event type (Table S2Q). Here again, the largest overlap was found for *SRSF3-high* and *SRSF7-high* tumors which shared 22% of their DSGs, whereas tumors with the most frequently altered SFs, *SRSF1-high, TRA2β-high* and *SRSF2-high* tumors shared between 10-13% of their DSGs (Table S2Q). Thus, although varying degrees of similarity and differences in splicing changes are detected, the overall splicing patterns are different in tumors with distinct SF alterations.

The finding that SFs are frequently altered in human breast tumors and are associated with the expression of distinct spliced isoforms prompted us to examine which of these SF alterations are likely to play a role in tumorigenesis. Therefore, we took advantage of an *in vitro* organotypic culture model to dissect the functional consequences of overexpression of each individual SF (SF-OE) on mammary transformation.

### Specificity of splicing factors in mammary cell transformation

We selected eight SFs upregulated in human breast tumors to systematically determine their individual roles in breast cancer initiation. Starting with the non-transformed human mammary epithelial MCF-10A cell line, we generated eight derivative cell lines, each stably overexpressing one of the SFs, using a retroviral construct containing a T7-tagged SF cDNA or an empty vector control (Figure 2A,B). Expression was confirmed by RT-qPCR, western blotting (Figure S3A,B), and immunofluorescence using a T7-tag antibody (Figure 2B). Cells overexpressing the oncogenic SF SRSF1 were previously shown to undergo transformation in this model (Anczuków et al., 2012) and were used as a positive control. As shown by immunofluorescence, all T7-tagged SFs were localized primarily to the nucleus, as expected (Figure 2B), consistent with their previously described role in splicing regulation (Cáceres et al., 1997). MCF-10A cells form organized growth-arrested three-dimensional (3D) structures when grown in Matrigel, a basement-membrane-like extracellular matrix, recapitulating the acinar structure observed in the mammary gland (Debnath and Brugge, 2005). Oncogenes associated with breast cancer were shown to disrupt the highly organized architecture of MCF-10A 3D acinar structures (Debnath and Brugge, 2005). We determined the effect of SF-OE on acinar morphology and size at days 8 and 16, and classified SFs into three categories, according to their effects on the acinar phenotype (Figure 2C,D): i) SRSF4-, SRSF6- and TRA2β-OE promoted the formation of dysmorphic acini that exhibited a 3-4 fold increase in acinar size *vs.* the empty vector control (*P*<0.0001); ii) HNRNPA1-OE, similarly to SRSF1 (Anczuków et al., 2012), increased acinar size by two-fold (*P* <0.001), but did not disrupt acinar morphology; iii) finally, the phenotypes of SRSF2-, SRSF3-, and SRSF9-OE were not significantly different from that of the empty-vector control (Figure 2C,D). Moreover, SRSF6- and TRA2β-OE-associated increases in acinar size were detected very early during acinar morphogenesis (Figure S4A,B), and were accompanied by increased numbers of proliferating day-8 acini (Figure S4C). Overall, our findings reveal that: i) not all SFs are oncogenic in this model; ii) only a subset of SFs are able to promote transformation of 3D-grown human mammary epithelial cells; and iii) distinct SFs promote different steps of tumorigenesis, suggesting underlying specificities and non-redundant functions of SFs, mediated by their downstream targets.

**Figure 2.**
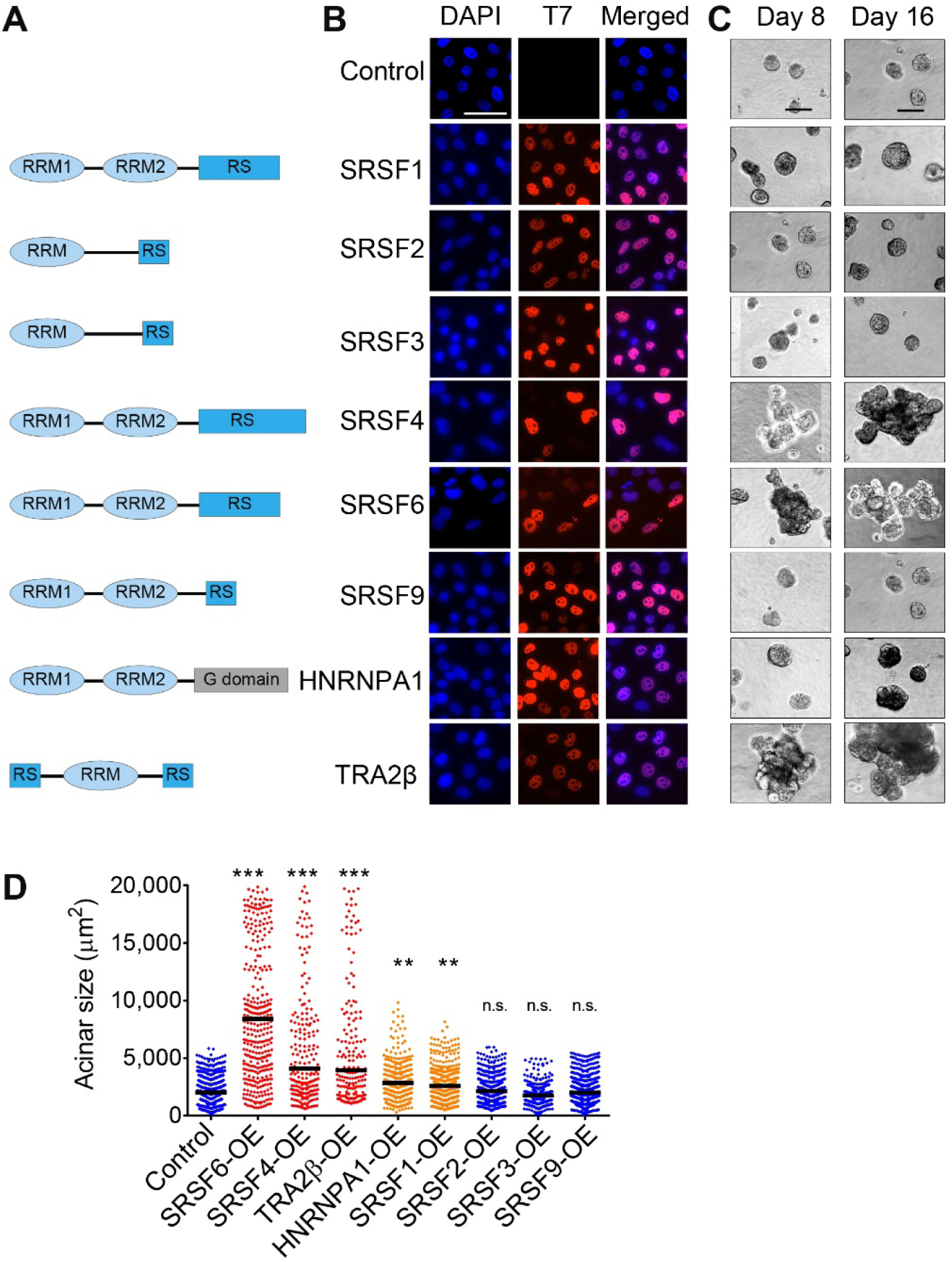
Specificity of splicing-factor-mediated transformation. **(A)** Graphical representation of the structural domains, *e.g.*, RNA-recognition motifs (RRM), serine/arginine-rich (SR) and glycine-rich (G)-domains, of the corresponding SFs shown in B. **(B)** Expression of T7-tagged SFs in MCF-10A cells, detected by immunofluorescence using T7-tag antibody and DAPI nuclei co-stain (scale bar: 50 μm). **(C)** Representative bright-field images of acinar size and morphology for control and SF-OE 3D MCF- 10A cells on days 8 and 16 (scale bar: 100 μm). **(D)** Quantification of acinar sizes in control and SF-OE 3D MCF-10A cells on day 16. The dot plot shows the size distribution of all structures and the median (horizontal line) for each condition (n=4, >50 acini per experiment; t-test \*\*\**P* <0.0001; \*\**P* <0.001; n.s. not significant compared to control).

### SRSF4-, SRSF6- and TRA2β-regulated alternatively spliced isoforms in breast cancer

Given the shared phenotype of SRSF4-, SRSF6-, and TRA2β-OE acini, we further investigated how these SFs contribute to transformation, by characterizing the DSEs each one of them regulates in model systems and human tumors. Very little is known about the splicing targets of these SFs; their function as splicing regulators has been investigated primarily in studies using minigene reporters, and also in several transcriptomic studies defining their endogenous targets (Long and Caceres, 2009), but no studies to date had used a cell context relevant to the mammary gland (Bradley et al., 2015; Jensen et al., 2014; Storbeck et al., 2014). To define the specificity and redundancy of these SFs, we performed RNA-seq on control and SF-OE MCF-10A day-8 acini and identified DSEs in: i) highly proliferative and dysmorphic SRSF4-, SRSF6-, and TRA2β-OE acini; and ii) SRSF2- and SRSF9-OE acini with no distinct phenotype *vs.* control. SRSF1 splicing targets were previously characterized by RNA-seq in MCF-10A cells using the same experimental strategy (Anczuków et al., 2015) and thus were not further investigated here.

Stranded Illumina poly-A selected RNA-seq libraries were generated for three biological replicates from day-8 acini and sequenced at >80 million reads per sample (Figure 3). Differential splicing analysis was performed as described above, in SF-OE *vs.* empty vector control MCF-10A. We identified >1500 DSEs in each condition, detected across multiple biological replicates (Figure 3A and Table S3A-E). Almost half of these DSEs correspond to CA events (Figure 3A). RI were the second most common class of DSEs, followed by alternative 5’ SS (A5’SS), alternative 3’ SS (A3’SS) and mutually exclusive exons (MXE) (Figure 3A). SRSF4-, SRSF6, and TRA2β-OE acini, which exhibit the strongest phenotypic changes compared to the control, displayed the largest number of DSEs (∼3,000-4000 events each; Figure 3A). Conversely, SRSF2- and SRSF9-OE acini, which were not phenotypically distinct from the control acini, affected ∼1500-2000 DSEs (Figure 3A). All five SFs promoted both inclusion (ΔPSI≥10%) and skipping (ΔPSI≤-10%) (Figure 3B), suggesting a dual role for SR proteins as splicing activators and repressors, either directly through RNA binding or indirectly through secondary interactions, consistent with previous findings in mouse, fruit fly, and human (Bradley et al., 2015; Pandit et al., 2013). A more detailed analysis by splicing-event type revealed that all five SFs promoted relatively equal numbers of inclusion and skipping CA events (Figure S5A). SRSF2-, SRSF4-, SRSF6-, and SRSF9-OE primarily promoted RI events, whereas TRA2β-OE promoted an equal number of retained and skipped introns (Figure S5A). For all five SFs, the majority of A5’SS and A3’SS events promoted usage of the proximal splice site (ΔPSI≥10%) (Figure S5A). To further understand the biological function of SFs in mammary epithelial transformation, we performed functional enrichment analysis of SF-regulated spliced isoforms in MCF-10A acini. Gene-set enrichment analysis revealed that SRSF4-, SRSF6-, and TRA2β-OE acini affect splicing of genes that regulate key cellular processes associated with transformation, such as the P53 pathway, DNA repair, G2M checkpoint, MYC target genes, E2F target genes, mitotic spindle, apical junctions, apoptosis, and metabolism (Figure S5B).

**Figure 3.**
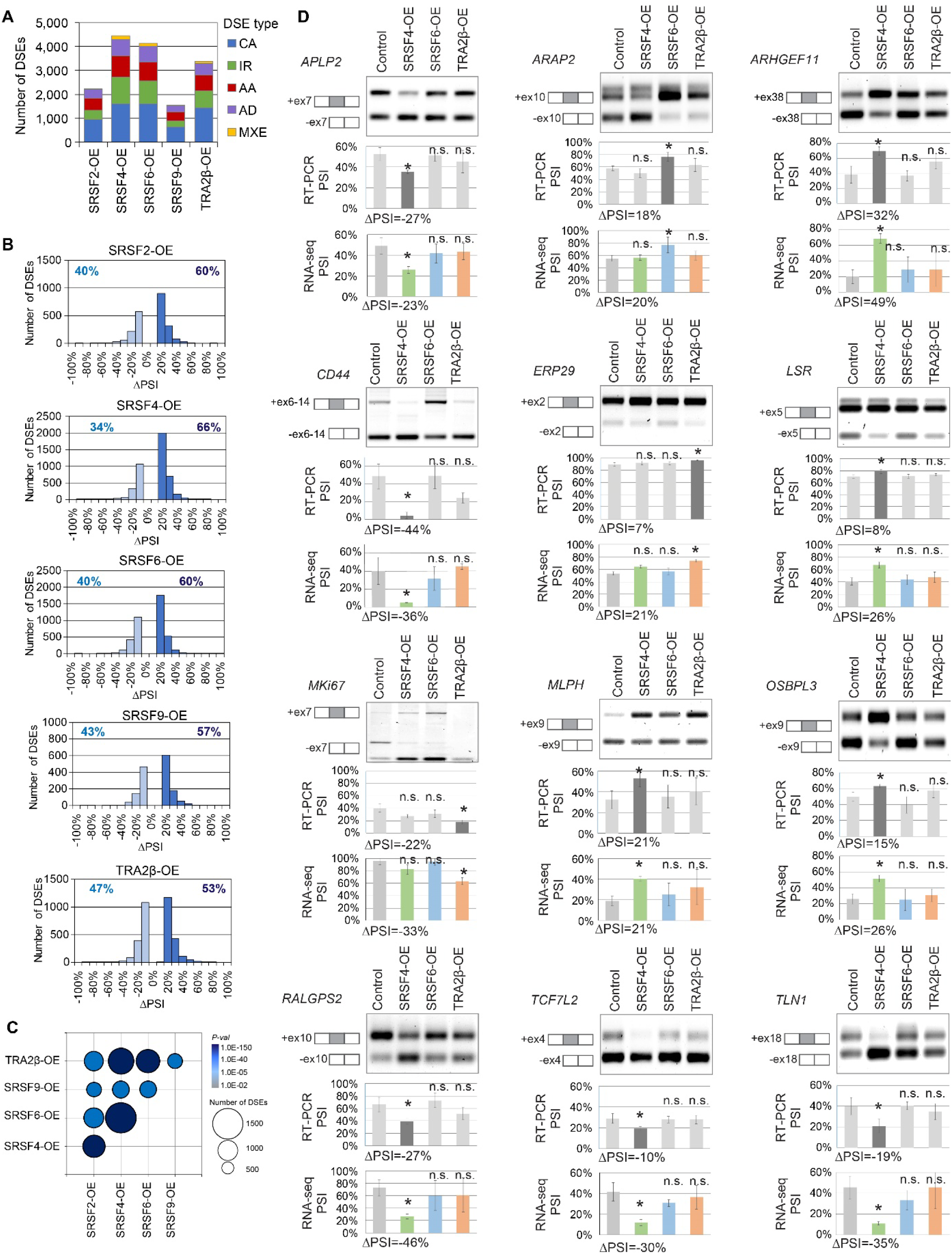
Splicing-factor overexpression is associated with differentially spliced events in MCF- 10A. **(A)** Number of differentially splicing events (DSEs) detected by RNA-seq in SF-OE *vs.* control MCF-10A day-8 acini (n=3; |ΔPSI|≤10%, FDR<5%, *P*<0.01), sorted by splicing event types (CA: cassette exon; MXE: mutually exclusive exon; RI: retained intron; A5’SS: alternative 5’ splice site; A3’SS: alternative 3’ splice site). **(B)** Skipped (ΔPSI≤-10%; light blue) and included (ΔPSI≥10%; dark blue) DSEs in SF-OE *vs.* control MCF-10A day-8 acini are plotted by ΔPSI values for each SF for all alternative splicing-event types (see Figure S5 for specific event types). The percentages of skipped and included DSEs are indicated for each SF. **(C)** Overlap in splicing targets in SF-OE MCF-10A day-8 acini for each indicated SF pair. The bubble size is proportional to the number of shared splicing events (DSEs); the color indicates the Fisher’s exact test P-value. See Table S3F and Methods for details. **(D)** RT-PCR validations of selected DSEs detected in SRSF4-, SRFS6-, or TRA2β-OE *vs.* empty vector control MCF-10A cells. A representative gel for each DSE is shown, along with the schematic isoform structure on the left. PSI values calculated from semi-quantitative RT-PCR (n=3; mean±SD; t-test, \**P*<0.05, n.s. not significant) and RNA-seq are plotted below each gel. Corresponding ΔPSI values are indicated below each graph for each significant DSE. See also Table S3 for details.

We then examined the overlap in splicing across all SF-OE MCF-10A datasets, either at the splicing-event level (DSEs) (Table S3F) or at the gene level (DSGs) (Table S3G). SRSF4-, SRF6-, and TRA2β- OE acini, which had a similar phenotype in 3D, also exhibited the greatest extent of overlap for both DSEs and DSGs, with 30-40% shared DSEs (Figure 3C and Figure S5B, Table S3F-G).

Next, we compared the SF-driven DSEs identified in MCF-10A acini to the DSEs detected in *SF-high* TCGA breast tumors (Figure S2B and Table S2B-O) to identify isoforms that likely contribute to SRSF4-, SRSF6-, or TRA2β-mediated transformation *in vitro* and are also detected in *SF-high* tumors. A total of 173 DSEs were detected in both SRSF4-OE MCF-10A acini and *SRSF4-high* TCGA tumors, 96 DSEs shared by SRSF6-OE MCF-10A acini and *SRSF6-high* TCGA tumors, and 36 DSEs shared by TRA2β- OE MCF-10A acini and *TRA2β-high* TCGA tumors (Table S3I). Almost 40% these DSEs affected regions with an annotated protein domain (Table S3I), suggesting that many of these splicing alterations could lead to the production of novel protein isoforms in breast tumors.

We validated 12 SF-regulated DSEs by semi-quantitative RT-PCR in SRSF4-, SRSF6-, and TRA2β-OE MCF-10A (Figure 3D). These splicing events impact genes previously shown to be involved in cell proliferation *(e.g., MKI67, APLP2*), cell migration and adhesion (*e.g., ARAP2, CD44, ERP29, LSR*), cell signaling (*e.g., ARHGEF11, MLPH, TCF7L2*) or cytoskeleton organization (*e.g., OSBPL3, RALGPS2, TLN1)*.

Since alternative splicing changes that affect protein domains have been shown to be critical for protein function in cancer (Climente-Gonzalez et al., 2017), we focused our analysis on DSEs that overlap with protein domains and are therefore likely to contribute to oncogenic phenotypes. Consistent with this idea, we observed that DSEs co-occurring in SRSF4-, SRSF6-, and/or TRA2B-OE significantly overlapped with annotated protein domains, compared to events unique to every SF (Figure S5D). In addition, to prioritize breast-cancer-relevant biological processes, we selected genes previously associated with breast cancer, as described in Open Targets (https://www.opentargets.org/), a database that uses human genetics and genomics data to prioritize drug targets for various diseases (Carvalho-Silva et al., 2019). We performed gene-enrichment analysis using enrichR (Chen et al., 2013a) on the subset of breast-cancer-associated genes with alternatively spliced protein domains. Analysis of each independent SF revealed that SRSF4-, SRSF6-, and TRA2β–OE targets, but not SRSF2- and SRSF9-OE targets, were significantly enriched in multiple cancer-relevant biological processes (Table S3K), suggesting that only the first three SFs impact mammary epithelial transformation through protein-altering AS. DSEs detected in all SRSF4-, SRSF6-, and TRA2β-OE acini were enriched in genes associated with the regulation of the cell cycle and G2/M transition, apoptotic processes, and Golgi-vesicle transport, as well as regulation of focal-adhesion assembly or regulation of cell-matrix adhesion (Table S3K). Altogether these results suggest a pivotal role of SRSF4, SRSF6, and TRA2β in the splicing-regulatory network that promotes mammary epithelial transformation, depending at least in part on discrete and effective coregulatory interactions with each other.

### Splicing-factor-regulated pathways in breast cancer

We also examined SF-induced gene-expression changes in MCF-10A acini using Cuffdiff (Trapnell et al., 2012) (Table S4A-F). SRSF4-, SRSF6-, and TRA2β-OE acini exhibited the greatest number of gene-expression changes, with >500 differentially expressed genes, respectively, *vs.* control acini (Figure S6A and Table S4B,C,E). In contrast to these SF-OE acini with high levels of expression changes, SRSF2- and SRFS9-OE acini exhibit only ≤40 genes with expression changes (Table S4A,D). Gene-set enrichment analysis suggested that genes altered in SRSF4-, SRF6-, or TRA2β-OE acini were enriched in targets associated with cell division, cell proliferation, cell-cycle control, cell migration, cytoskeleton organization, the extracellular matrix, cell polarity, cell signaling, or cholesterol metabolism (Table S4H-J). In addition, SRSF4-, SRSF6-, and TRA2β-OE acini shared 15-28% of differentially expressed genes (Figure S6B and Table S4F). The differentially expressed genes shared by SRSF4-, SRSF6-, and TRA2β-OE acini were associated with cell proliferation and the cell cycle, cytokine-mediated signaling, cholesterol biosynthesis, cell migration, and extracellular matrix organization (Table S4K-M and Figure S6C). Thus, SRSF4, SRSF6, and TRA2β share not only a similar phenotype in MCF-10A cells, but also regulate the expression of shared genes and spliced isoforms that affect cellular processes associated with tumor initiation and progression.

### TRA2β cooperates with the MYC breast-cancer oncogene

Transformation often results from cooperation among several oncogenes (Pedraza-Farina, 2006). We previously demonstrated that in breast cancer SRSF1 cooperates with MYC (Anczuków et al., 2012), one of the most frequently altered genes in cancer; yet, it remains unknown if this cooperation is specific to SRSF1 or whether it extends to other SR proteins. Here, we generated MCF-10A cells overexpressing selected SFs together with an inducible MYC transgene (Eilers et al., 1989) and assessed their effects on acinar morphogenesis. TRA2β-OE together with MYC promoted the formation of acinar structures that were enlarged compared to those resulting from use of MYC alone or TRA2β alone (Figure 4A, B). In contrast, no significant differences in size or morphology were observed following overexpression of SRSF6 or SRSF4 together with MYC (Figure 4B). Furthermore, we showed that *MYC* and *TRA2β* are co-expressed at high levels in human breast tumors (*P*<2.55E-10), as well as in several other tumor types (Figure 4C), suggesting that this cooperation also happens *in vivo*. Additionally, we determined that *TRA2β* mRNA expression increased 4 hours after MYC induction in MCF-10A cells, and increased protein levels are detected at 24h (Figure 4D), suggesting that *TRA2β* is likely a direct transcriptional target of MYC. Direct binding of MYC to the *TRA2β* proximal promoter region is further supported by genome-wide chromatin immunoprecipitation (ChIP)-sequencing data from mammary MCF-7 and MCF-10A cells from the ENCODE database (Figure 4E). Finally, MYC is amplified and/or upregulated in 51% of *TRA2β-high* breast tumors *vs.* 26% of *TRA2β-low* tumors (Figure 4F) (*P*<2.9E-05). Thus, our findings suggest that *TRA2β*-OE might be driven by MYC alterations in at least a subset of breast tumors.

**Figure 4.**
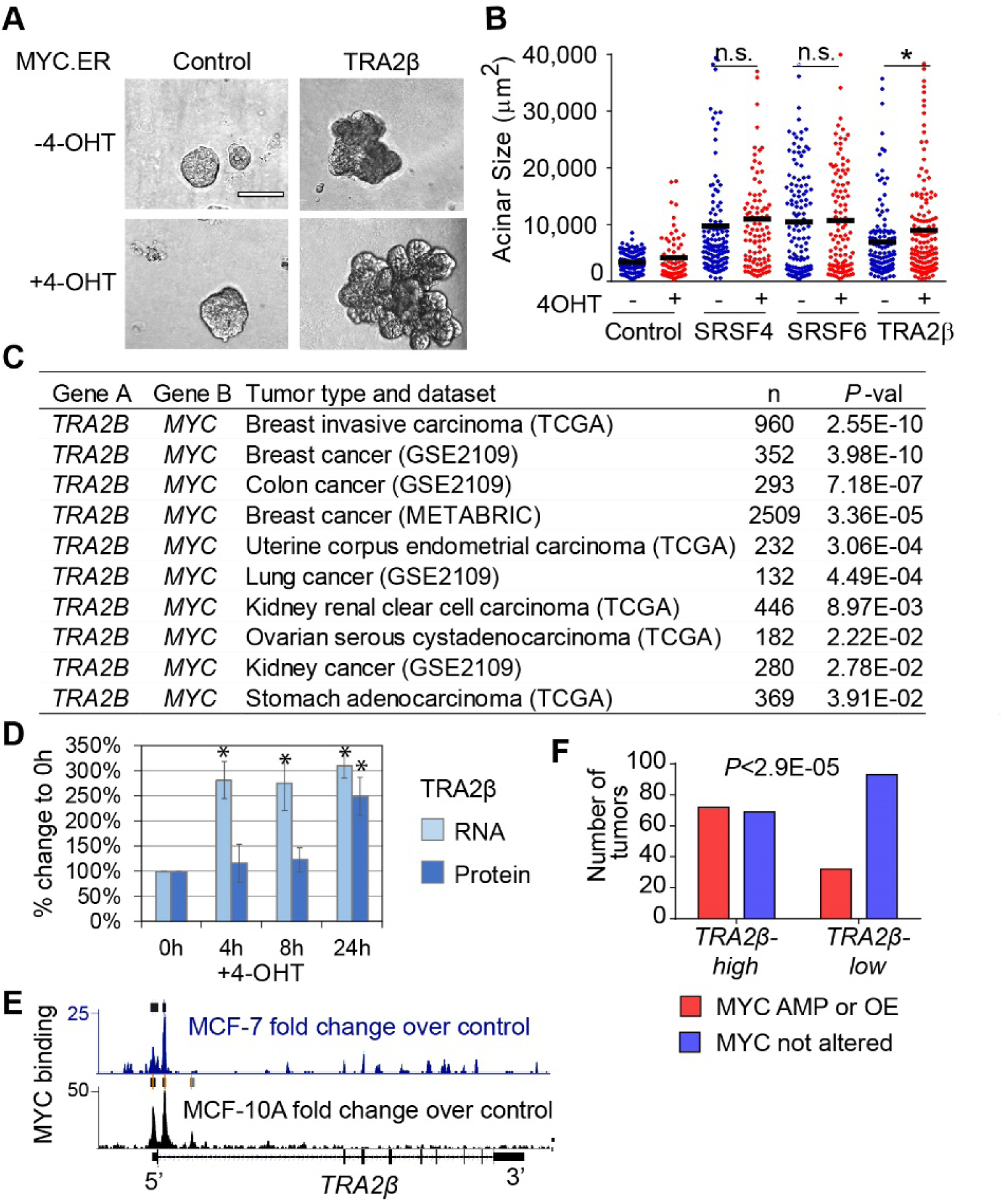
Cooperation of TRA2β with the MYC oncogene in breast cancer. **(A)** Representative bright-field images of empty-vector control or TRA2β-OE 3D MCF-10A acini expressing the estrogen-receptor (ER)-inducible MYC (Eilers et al., 1989), which is activated by 4-OHT treatment (scale bar: 100 μm). **(B)** Acinar size distribution of SRSF4-, SRSF6-, TRA2β-OE MCF-10A cells expressing MYC.ER +/- 4- OHT (n≥3, >100 acini per experiment; t-test * *P*<0.01, n.s. not significant; median is shown as horizontal line). **(C)** Correlation between *MYC* and *TRA2β* expression in human tumors. Dataset IDs and numbers of samples are indicated. *MYC* and *TRA2β* expression were grouped in four categories: i) both low; ii) both high; iii) low *MYC* and high *TRA2β*; and iv) high *MYC* and low *TRA2β. P*-values from the Fisher exact test are shown. **(D)** Changes in TRA2β RNA and protein levels in MCF-10A MYC.ER following MYC activation with 4- OHT at the indicated time points were detected by qRT-PCR and western blotting respectively (n=3; mean±SD; t-test, \**P*<0.05). RNA levels are normalized to GAPDH and HPRT, protein levels are normalized to Tubulin. **(E)** MYC binding to *TRA2β* genomic region as detected by MYC ChIP-seq experiments in MCF-7 (ENCSR000DMJ) and MCF-10A (ENCSR000DOS) cells. Tracks show fold changes over control, as calculated from ENCODE for pooled replicates for each cell line. Significant peaks called using conservative IDR thresholds are shown by rectangles above each track. A schematic representation of the *TRA2β* gene is shown below, in the 5’ to 3’ orientation. **(F)** MYC amplification (AMP) or MYC overexpression (OE) (Z-score ≥2) status shown for TCGA *TRA2β- high* (Z-score ≥2) *vs. TRA2β-low* (Z-score <0) breast tumors. *P*-value from the Fisher exact test is indicated.

### Differential role of splicing factors in cell migration and invasion

The highly dysmorphic phenotype of SRFS4-, SRSF6-, and TRA2β-OE acini (Figure 2C,D), combined with the splicing and gene-expression changes detected by RNA-seq in targets associated with cell migration and cytoskeleton organization (Tables S3 and S4), suggested that these SFs could play a role in cell migration and invasion. To assess how SF-OE affects different types of cell movement, we performed a series of 2D and 3D cell-based assays in non-invasive MCF-10A cells (Figure 5 and S7). In 3D collagen-matrigel assays (Xiang and Muthuswamy, 2006), the formation of intra-acinar bridges and multicellular protrusions, which are indicative of invasive behavior, was observed in ∼25-50% of the SRSF4-, TRA2β-, HNRNPA1-, and SRSF6-OE acini, but in only <1% of control acini (Figure 5A,B). Furthermore SRSF4-, TRA2β-, and SRSF6-OE increased cell movement within acini structures, as detected by live-cell imaging (Figure S7A), and SRSF6-OE also significantly increased cell migration in 2D transwell assays (Figure S7C). Surprisingly, despite the lack of effect of SRSF9-OE on acinar morphology or size, SRSF9-OE cells migrated faster both in wound-healing and transwell-migration assays, compared to the control (Figure S7B,C), and exhibited invasive behavior in 3D collagen-matrigel assays (Figure 5A,B). These results suggest that the effects of SF-OE on cell proliferation and cell invasion can be uncoupled.

**Figure 5.**
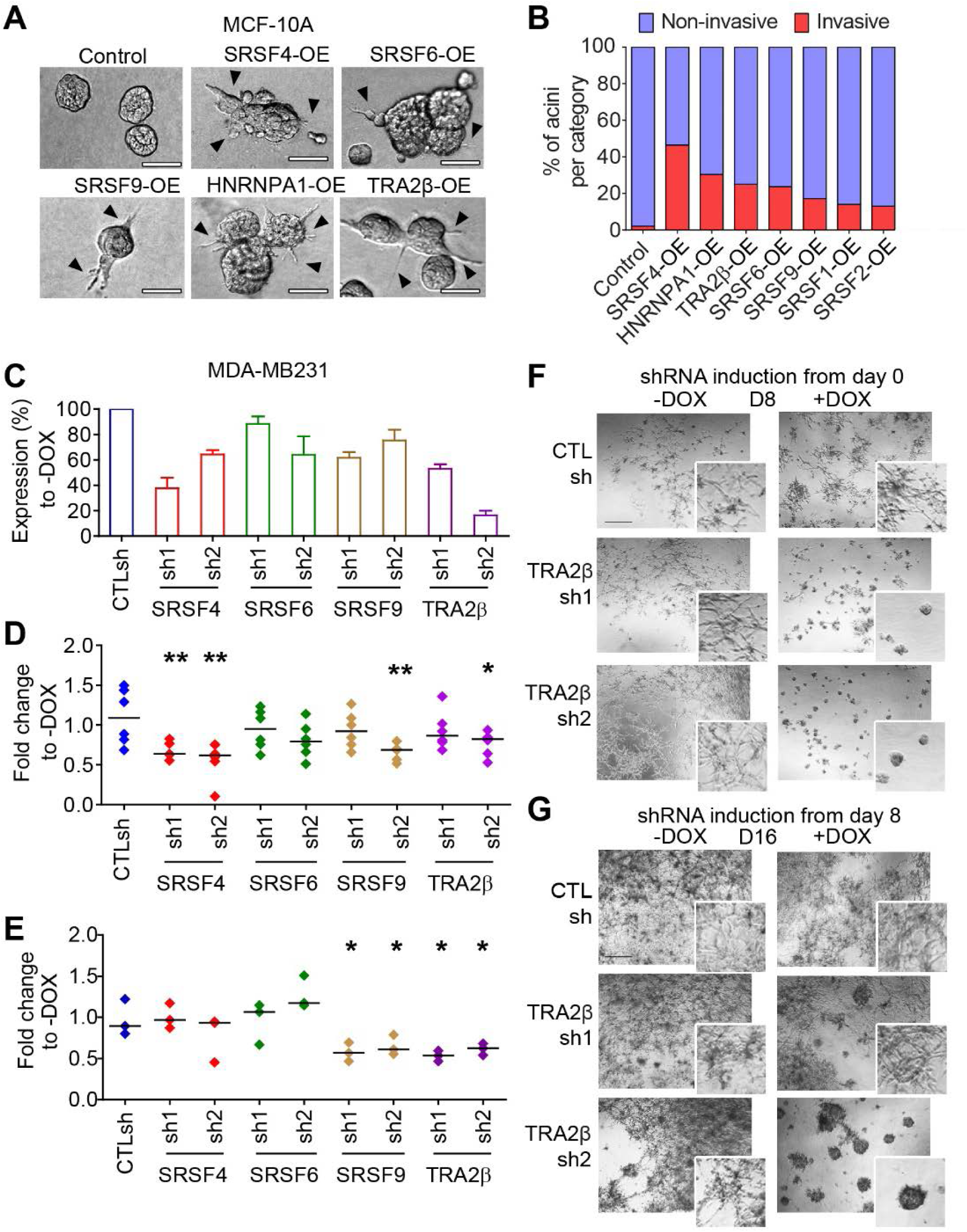
Differential roles of splicing factors in cell migration and invasion. **(A)** Representative bright-field images of acinar morphology for control or SF-OE 3D MCF-10A cells grown in matrigel-collagen invasion assay at day 8. Multicellular protrusions and inter-acinar bridges representative of invasive structures are indicated by arrowheads (scale bar: 50 μm). **(B)** Percent of invasive *vs.* non-invasive acini in SF-OE 3D *vs.* control MCF-10A cells imaged at day 8 as in A (n=3, >50 acini per condition). **(C)** SF expression in MDA-MB231 cells stably expressing doxycycline (DOX)-inducible SF-targeting or control (CTLsh) shRNAs. SF levels were quantified 72 hours after DOX treatment by western blotting using SF-specific antibodies, normalized to a tubulin loading control. The % of SF expression for each shRNA +DOX is normalized to the corresponding -DOX. (n=3; mean±SD). **(D-E)** Migration of MDA-MB231 cells expressing a control or SF-targeting shRNA in 2D wound-healing or 2D transwell assay (E). The plot shows the distribution and the median (horizontal line) for each condition +DOX normalized to the corresponding -DOX (n=3; t-test, \*\**P*<0.005, \**P*<0.05). **(F-G)** Representative bright-field images of MDA-MB231 cells grown in 3D in matrigel expressing either a control (CTLsh) or TRA2β-targeting shRNA (TRA2βsh1 or sh2) (scale bar: 100 μm). shRNAs are induced by adding DOX either from day 0, when the cells are seeded (F), or from day 8, after cells have formed invasive structures (G).

We then assessed whether SF expression is required to maintain the migratory phenotype of metastatic breast-cancer cells. We focused on a TNBC model, as TRA2β is upregulated in 40% of TNBC tumors (Figure S1C), an aggressive cancer subtype associated with high metastasis incidence. We examined how SRSF4, SRSF6, SRSF9, and TRA2β—which increased migration and invasion in MCF-10A cells when overexpressed—affected the invasive potential of human MDA-MB231 TNBC cells. We established stable cell lines with a doxycycline (DOX)-inducible short hairpin RNA (shRNA) expression system that enables precise tracking of retroviral transduction and shRNA induction through two fluorescent reporters, *i.e.*, constitutively expressed GFP as well as DsRed that is expressed along with the shRNA (Zuber et al., 2011). Six inducible shRNAs were tested for each SF; then, for each SF, the two shRNAs that promoted the strongest knockdown (KD), along with the control scrambled shRNA (CTLsh), were selected for further assays (Figure 5C). Importantly, SF-KD did not decrease cell-doubling time in MDA-MB231 cells (Figure S8A,B), allowing us to define the consequences of SF-KD on cell invasion, independently of its potential effect on cell proliferation. TRA2β-KD and SRSF9-KD both decreased cell migration in wound-healing and transwell-migration assays (Figure 5D,E). A milder phenotype was observed for SRSF4-KD, which decreased migration only in wound-healing assays (Figure 5D). Finally, we assessed SF-KD in MDA-MB231 cells grown in 3D culture in matrigel, where these transformed cells form stellate structures, representative of their invasive behavior (Kenny et al., 2007). When the shRNAs were induced from day 0, TRA2β-KD, with either shRNA, prevented the formation of stellate invasive structures (Figure 5F and S8C-E), and promoted the formation of round ‘acinar-like’ structures lacking invasive behavior (Figure S8E). SRSF6-KD also prevented the formation of invasive structures, whereas SRSF4- and SRSF9-KD decreased the number of stellate structures, as well as their size (Figure S8C-D). Finally, taking advantage of the inducible system, we asked whether TRA2β is required to maintain the invasive phenotype after the structures have already formed. Inducing the TRA2β-targeting shRNA on day 8 after the formation of invasive structures was sufficient to reverse the phenotype and promote the formation of non-invasive acinar-like structures (Figure 5G and S8E). The effect seemed to be dose-dependent, as the weaker shRNA, TRA2βsh1, had an intermediate phenotype compared to that of the stronger shRNA TRA2βsh2 (Figure 5G). In summary, our findings suggest that specific SFs play a role in controlling cell migration during the early steps of transformation, or in maintaining the invasive potential in established breast-cancer cell lines. In particular, TRA2β expression was sufficient to initiate invasion in non-transformed mammary epithelial cells, and was required to maintain TNBC cell invasion.

### TRA2β targets in breast-cancer metastasis

To define the TRA2β-regulated isoforms that play a role in TNBC cell invasion, we performed RNA-seq and identified DSEs by comparing day-8 3D-grown MDA-MB231 cells in three conditions: i) TRA2βsh1+DOX *vs.* -DOX; ii) TRA2βsh2+DOX *vs.* -DOX; and iii) CTLsh+DOX *vs.* -DOX. Stranded Illumina poly-A selected RNA-seq libraries were generated for three biological replicates, and sequenced at >90 million reads per sample. DSEs were identified as described above. 3D-grown MDA-MB231 cells expressing CTLsh+DOX *vs.* -DOX were used to identify DSEs induced by DOX treatment, *i.e*., the significant DSEs from TRA2βsh+DOX *vs*. -DOX were discarded if they also appeared as significant and altered in the same direction in CTLsh+DOX *vs.* -DOX (Table S5A). Following exclusion of the DOX-induced DSEs, we identified >3,500 DSEs in TRA2βsh2+DOX *vs.* -DOX, and >2,000 in TRA2βsh1+DOX *vs.* -DOX (Figure 6A and Table S5B,C); the number of DSEs associated with the stronger shRNA TRA2βsh2 was higher than the number associated with the weaker shRNA TRA2βsh1. TRA2βsh1+DOX and TRA2βsh2+DOX cells shared 582 DSEs (*P*=1.5E-35), as well as 691 DSGs (*P*=3.3E-14) (Table S5E-F). In MDA-MB231 cells, TRA2β-KD promoted primarily CA events, and promoted more skipping than inclusion (Figure 6A,B). Gene-set enrichment analysis revealed that TRA2βsh1+DOX and TRA2βsh2+DOX cells affected splicing of genes that regulate key cellular processes associated with transformation, such as the P53 pathway, DNA repair, G2M checkpoint, MYC target genes, E2F target genes, mitotic spindle, apical junctions, apoptosis, and metabolism (Figure 6C). Analysis of breast-cancer-associated genes with alternatively spliced protein domains revealed that TRA2β-regulated targets were enriched in genes associated with G2/M regulation, protein deubiquitination, DNA repair, regulation of cell migration, and regulation of apoptotic process (Table S5G-H). Interestingly, these biological processes were also highly ranked in the TRA2β splicing targets in MCF-10A cells, albeit represented by different set of genes (Table S5H). These observations presumably reflect differences in spliced isoforms associated with TRA2β function in tumor initiation in non-transformed MCF-10A *vs.* its role in tumor maintenance and invasion in metastatic MDA-MB231 cells.

**Figure 6.**
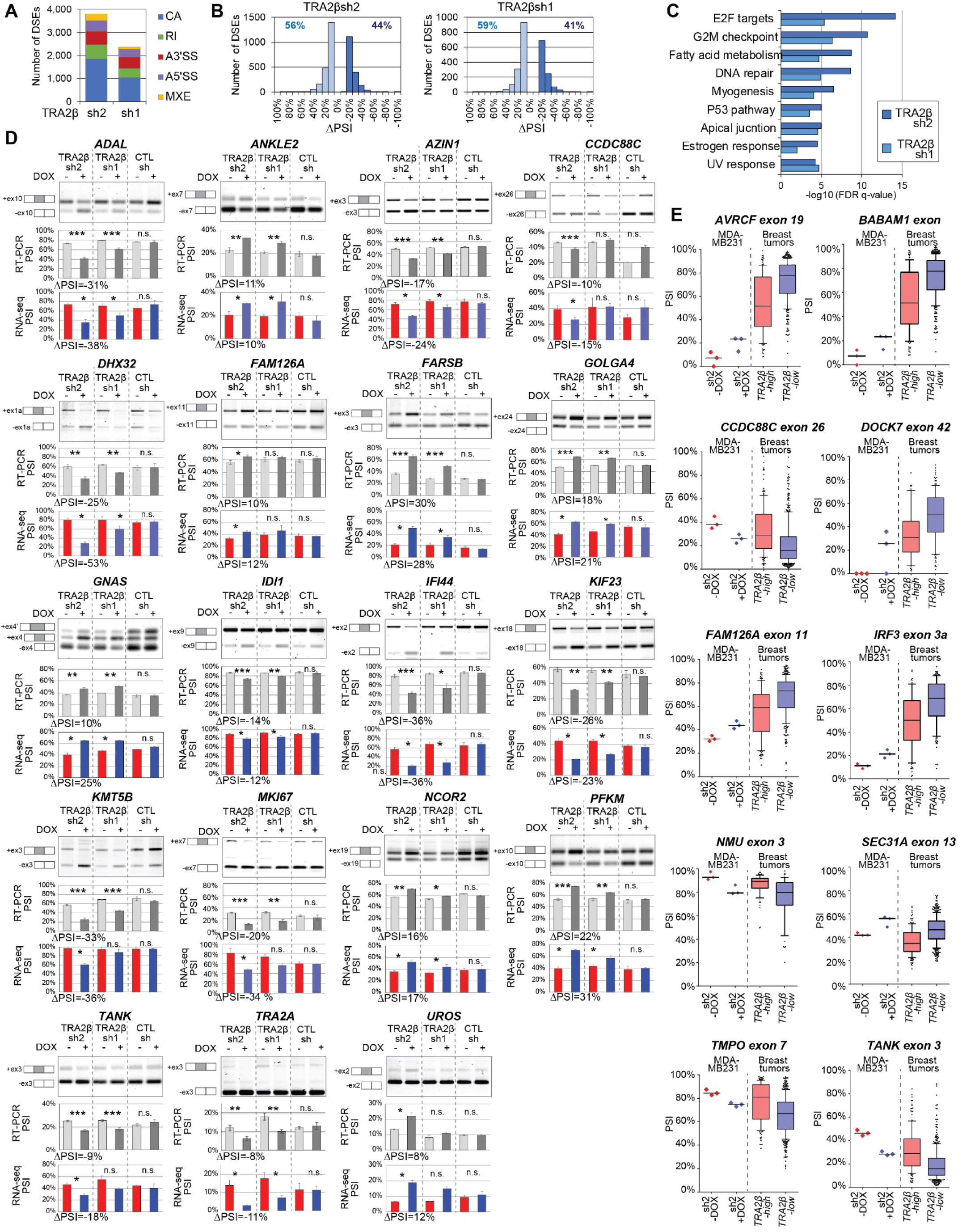
TRA2β-KD promotes changes in spliced isoforms in 3D MDA-MB231 cells. **(A)** Number of differentially splicing events (DSEs) detected by RNA-seq in TRA2βsh2 +DOX *vs.* -DOX, and TRA2βsh1+DOX vs. -DOX, in 3D MDA-MB231 cells at day 8 (n=3; |ΔPSI|≥10%, FDR<5%, *P*<0.01), sorted by splicing event types (CA: cassette exon; MXE: mutually exclusive exon; RI: retained intron; A5’SS: alternative 5’ splice site; A3’SS: alternative 3’ splice site). DOX-induced DSEs detected in CTLsh+DOX *vs.* -DOX were removed (see Methods). **(B)** Skipped (ΔPSI≤-10%; light blue) and included (ΔPSI≥10%; dark blue) DSEs in TRA2βsh2+DOX *vs.* -DOX, or TRA2βsh1+DOX *vs.* -DOX, are plotted by ΔPSI values. The percentage of skipped and included DSEs is indicated for each condition. **(C)** Gene Set Enrichment Analysis for DSEs detected in TRA2βsh2 or TRA2βsh1 3D MDA-MB231 cells at day 8. The top 10 Hallmark gene sets are shown. **(D)** RT-PCR validations of selected DSEs detected in TRA2βsh2 or TRA2βsh1 3D MDA-MB231 cells on day 8. A representative gel is shown, along with the schematic isoform structure on the left. Corresponding PSI values for all samples and ΔPSI for TRA2βsh2 calculated from semi-quantitative RT- PCR (n=3; mean±SD; t-test, \*\*\**P*<0.0005; \*\**P*<0.005; \**P*<0.05; n.s. not significant) and RNA-seq are plotted below each gel. See Table S5 for details. **(E)** PSI values of significant DSEs detected in *TRA2β-high* (n=146) vs. *TRA2β-low* (n=449) TCGA breast tumors and also detected in TRA2βsh2+DOX *vs.* -DOX MDA-MB231 cells (n=3). See Table S5H for details.

For many TRA2β-regulated targets, we detected a dose-dependent effect of the two shRNAs, *i.e.*, the splicing changes were stronger with TRA2βsh2 than with TRA2βsh1. We validated sixteen TRA2β-regulated DSEs by semi-quantitative RT-PCR in 3D-grown MDA-MB231 cells (Figure 6D). These splicing events impact genes previously shown to be involved in mitosis regulation (*ADAL*), cell migration and adhesion (*DHX32, IFI44, KMT5B, KIF23, TANK*), cell signaling (*FAM126A, CCDC88C, GNAS, NCOR2, TANK*), splicing (*TRA2α*) or cell metabolism (*ADAL, FARSB, IDI1, UROS, PFKM)*.

Additionally, we measured how TRA2β-KD affects gene expression in 3D MDA-MB231 cells (Table S6). We identified >1,400 upregulated and >1,200 downregulated genes in TRA2βsh2+DOX *vs*. -DOX (Table S6B), whereas the weaker hairpin, TRA2βsh1, had an intermediate effect (Table S6C). Genes affected by TRA2β-KD were associated with Wnt signaling, control of cell polarity, cell cycle regulation, and transcriptional control (Table S6D). Differential splicing also led to changes in gene expression; indeed, 312 genes were affected both at the splicing and gene-expression levels in TRA2βsh2+DOX (Table S6E). To further define which spliced isoforms relevant to human breast tumors, we focused on DSEs detected both in MDA-MB231 cells with high TRA2β levels as well as *TRA2β*-*high* breast tumors. We identified 32 shared DSEs, defined as events regulated in opposite directions, *i.e.*, ΔPSI≥10% in TRA2βsh2+DOX *vs*. -DOX MDA-MB231 and ΔPSI≤-10% in *TRA2β*-high *vs. TRA2β*-low tumors, and vice-versa (Table S5K and Figure 6E). These DSEs affect genes previously linked with cancer and/or metastasis, *e.g.*: i)skipping of exon 19 of *ARVCF*, a gene involved in cell adhesion, and often downregulated in breast tumors (Anastasiadis and Reynolds, 2000; Xiang et al., 2015). Exon 19 encodes the last 20 amino acids of the protein; ii) usage of an alternative 3’SS in exon 2 of *BABAM1*, a gene that encodes a member of the BRISC and BRCA1 complex (Feng et al., 2009; Shao et al., 2009); iii) inclusion of *CCDC88C* exon 26, a signal transducer involved in Wnt signaling that can fuel the epithelial to mesenchymal transition (EMT), trigger cancer-cell migration and invasion, and drive metastasis (Ara et al., 2016; Aznar et al., 2015; Ishida-Takagishi et al., 2012; Kobayashi et al., 2005). Inclusion of exon 26 introduces a PTC, thereby truncating the last 493 amino acids of the protein, and producing a transcript that could potentially be degraded by NMD; iv) skipping of exon 42 of *DOCK7*, a gene involved in cell invasion in both a Rac-dependent and Rac-independent manner (Murray et al., 2014; Yang et al., 2012). Exon 42 encodes 5 amino acids in the C-terminal DHR-2 catalytic domain and thus its skipping could affect DOCK7 activity; v) skipping of exon 11 of *FAM126A*, which encodes a protein involved in localizing PI4K to the plasma membrane and regulating phosphatidylinositol 4-phosphate synthesis (Baskin et al., 2016). Inclusion of exon 11 leads to a C-terminally truncated protein; vi) inclusion of exon 3a of *IRF3*, leading to the expression of an spliced isoform that differs by 20 amino acids in the N-terminal region (Karpova et al., 2000); vii) skipping of exon 3 in *NMU*, which encodes the NMUR2S spliced isoform that lacks one of the six transmembrane domain. Expression of NMUR2S blocks endogenous NMU signaling and suppresses proliferation in ovarian cancer cells (Lin et al., 2015); viii) inclusion of exon 13 of *SEC31A*, a spliced isoform previously associated with EMT (Xu et al., 2014). ix) inclusion of exon 7 in *TMPO*, a gene that encodes a protein involved in nuclear-structure organization and cell-cycle dynamics (Dechat et al., 2000). Exon 7 encodes a Lamin B binding domain, and exon-7-containing isoforms are enriched in human breast tumors (Marrero-Rodriguez et al., 2015). Moreover, KD of *TMPO*+ex7 inhibits cell proliferation and suppresses metastasis in pancreatic cancer (Kim et al., 2012); X) inclusion of exon 3 of *TANK*, a gene associated with cell migration and tumor necrosis factor receptor signaling. In glioblastoma, TANK-KD arrests cells in the S-phase and prohibits tumor cell migration (Stellzig et al., 2013). Overall our findings suggest that TRA2β regulates splicing of multiple genes associated with celladhesion, invasion, and metastasis, as well as cell-cycle regulation and DNA replication in both breast-cancer models and patients’ breast tumor tissues.

### TRA2β plays a role in breast-cancer metastasis *in vivo* and its expression correlates with clinical outcomes in breast-cancer patients

Finally, to assess the *in vivo* effect of TRA2β-KD on cell invasion and metastasis, TRA2βsh2 or CTLsh MDA-MB231 cells were orthotopically injected in the mammary gland of NOD.Cg-*Prkdc*^scid^ *Il2rg*^tm1Wjl^/SzJ (NSG) female mice (Figure 7A). Primary tumor and metastasis formation were monitored by luciferin bioluminescence imaging. TRA2β-KD did not prevent the formation of primary mammary tumors (Figure 7B and S9A-C) nor did it affect the size of primary tumors, compared to the non-induced (TRA2βsh2+DOX *vs.* –DOX) and induced controls (TRA2βsh2+DOX *vs.* CTLsh +DOX) (Figure S9B). However, we observed a decrease in the metastatic burden (*i.e.*, percent of metastatic organ area relative to the whole organ area) in both lung and liver in TRA2β-KD animals, compared to non-induced (TRA2βsh2+DOX *vs.* -DOX) and induced controls (TRA2βsh2+DOX *vs.* CTLsh+DOX) (Figure 7C,D), as well as decrease in lung, liver, or lymph node macro-metastases (Figure S9D). TRA2β-KD reduced the number of metastatic foci, but had no significant effect on focus size (Figure S9B-J). Additionally, we assessed the role of TRA2β levels in the latter steps of the metastatic cascade, *i.e.*, survival in blood circulation, extravasation, and colonization of a distant organ site (Lambert et al., 2017), by monitoring metastasis formation following tail-vein injection of cancer cells (Figure S10A). Here, TRA2β-KD (TRA2βsh2+DOX *vs.* -DOX) also reduced lung metastasis formation (Figure S10B-D). Finally, clinical data for human breast tumors revealed that *TRA2β*-*high* levels are associated with a decrease in overall survival (Figure 7E), as well as in distant-metastasis-free survival (Figure 7F) in multiple patient cohorts. Overall, our data strongly suggest a role for TRA2β and its splicing targets in regulation of breast-cancer metastasis initiation and maintenance.

**Figure 7.**
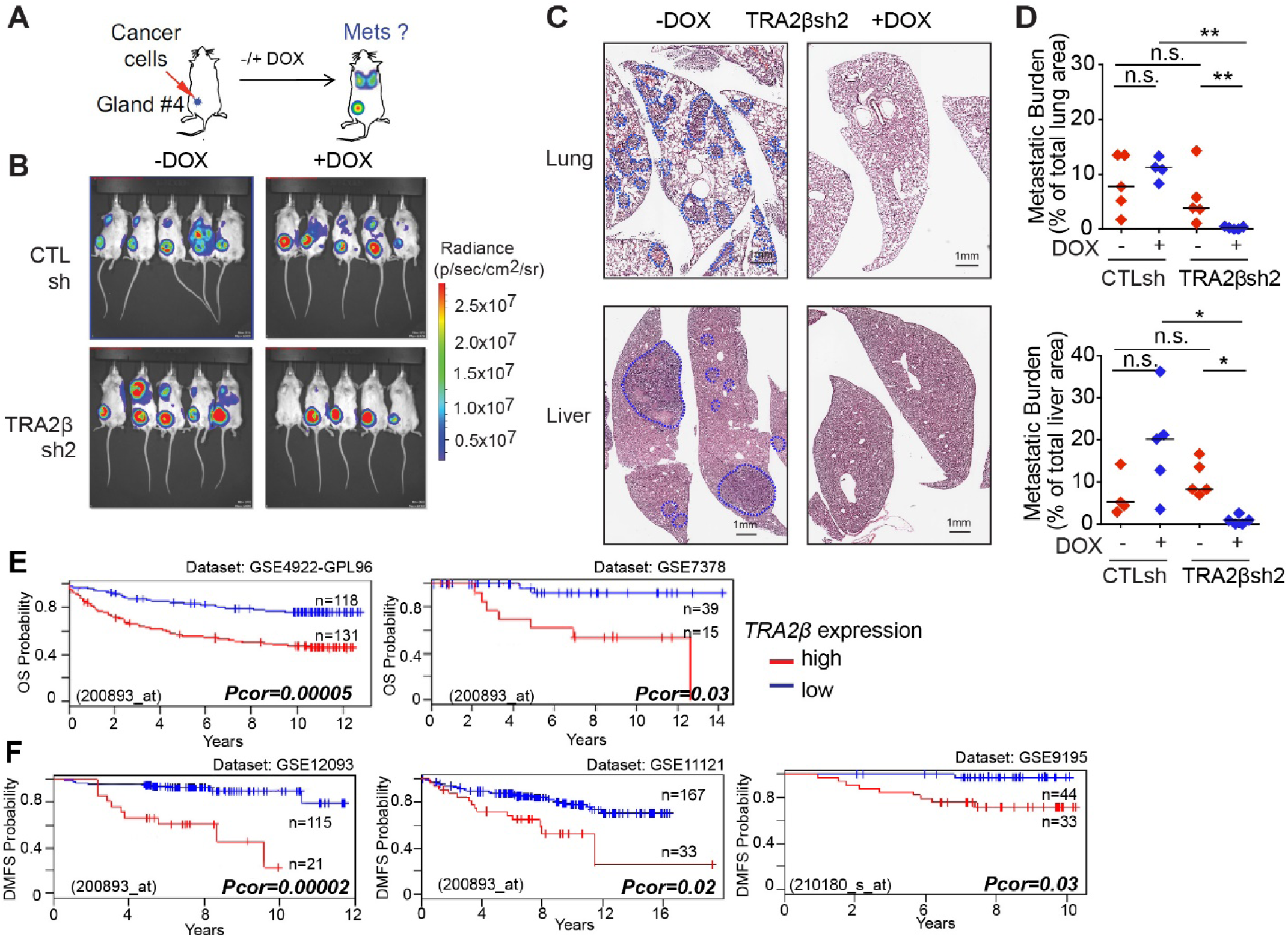
TRA2β plays a role in metastasis *in vivo* in a TNBC mouse model and in breast-cancer patients. **(A)** MDA-MB231 cells expressing CTLsh or TRA2βsh2 were injected into the mammary fat pad of NSG mice; primary tumors and metastasis were monitored by bioluminescence imaging and histopathology. **(B)** Bioluminescence detection of primary tumors and metastasis in mice injected with CTLsh or TRA2βsh2 MDA-MB231 cells +DOX *vs.* -DOX at 8 weeks post-injection. **(C)** Representative H&E pictures from lung and liver sections of mice injected with TRA2βsh2+DOX *vs.* -DOX MDA-MB231 cells (scale bar: 1 mm). Metastatic areas are circled in blue. Metastases are scored through the whole organ, using sections evenly distributed every 2 mm and quantified in D. **(D)** Quantification of metastasis burden, *i.e.*, percentage of metastatic organ area relative to the whole organ area, in mice injected with CTLsh or TRA2βsh2 MDA-MB231 cells +DOX *vs.* -DOX (n≥4; t-test, \*\**P*<0.001, \**P*<0.01, n.s. not significant). **(E-F)** Correlation between *TRA2β* expression and overall survival (OS) (E) or distant-metastasis-free-survival (DMFS) (F) in different cohorts of breast-cancer patients stratified by *TRA2β* levels. High (red) and low (blue) expression groups are dichotomized as described (Mizuno et al., 2009). Cohort size, GEO dataset and probe ID, and corrected *P*-value (log-rank test) for each dataset are indicated.

## DISCUSSION

### Splicing-factor alterations in mammary cells have distinct functional consequences

Abnormal alternative splicing has emerged as a hallmark of human tumors (Urbanski et al., 2018). Defects in splicing regulators are associated with many tumor types (Dvinge et al., 2016), yet for most SFs their functional role in tumorigenesis remains poorly characterized. Here, we systematically identify alterations in the SR protein family and their frequency across a large collection of human breast tumors. We then use breast cancer models to define the functional consequences of alterations in these SFs and identify their splicing targets, uncovering distinct functions for each SR protein in mammary tissues.

Our analysis of >900 human breast tumors revealed that SFs are frequently altered, but exhibit distinct alteration patterns and subtype specificity (Figure 1 and S1). In addition to the breast-cancer oncogeneSRSF1 (Anczuków et al., 2012), *TRA2β, SRSF2*, and *SRSF6* are each overexpressed ≥10% of tumors, all subtypes combined. Interestingly, *TRA2β-, SRSF12-, SRSF3-*, and *SRSF2-*high tumors represent >20-40% of TNBCs, the most aggressive breast-cancer subtype (Figure S1). We determined that tumors with high levels of a given SF express unique DSEs, compared to tumors with low levels of that SF, and that many DSEs are recurrently detected in >100 tumor samples (Figure S2 and Table S2). Additionally, tumors with alterations in distinct SFs share relatively low numbers of DSEs (<20% overlap), with the exception of *SRSF3*-high and *SRSF7*-high tumors, which share 46% of their DSEs and 22% of their DSGs (Table S2). Our findings thus suggest that distinct SFs have different targets and biological functions in breast tumors.

By overexpressing selected SFs in non-transformed mammary epithelial cells, we demonstrated that SRSF4-, SRSF6-, and TRA2β-OE promote transformation, as well as cell migration and invasion (Figure 2 and 5). Importantly, we revealed that not all SFs are oncogenic in this context. Indeed, although *SRSF2* is overexpressed in 13% of human breast tumors (Figure 1), SRSF2-OE did not affect any of the measured phenotypes in non-transformed mammary epithelial cells (Figure 2). Strikingly, recurrent mutations in *SRSF2* are found in human hematologic malignancies, but not in other tumor types (Visconte et al., 2012). Functionally, blood lineage-specific expression of *Srsf2*^*P95H*^ causes defective hematopoiesis (Kim et al., 2015), and *SRSF2* mutations alter its RNA-binding preference, thereby altering its target recognition in the hematopoietic lineage (Kim et al., 2015). Our findings demonstrate that changes in *SRSF2* levels trigger relatively few splicing alterations in mammary epithelial cells (Figure 3 and Table S3), and suggest that SRSF2 may not contribute to early steps of tumor initiation in the breast. Our study also suggests a restricted role for SRSF9 in tumorigenesis, which when overexpressed in mammary cells increased cell migration and invasion, but did not affect cell proliferation, apoptosis, or acinar architecture (Figure 1 and 5). Previous studies are consistent with a role for SRSF9 in cell invasion. In addition to breast, SRSF9 is also overexpressed in human melanoma, colon cancer, and glioblastoma (Fu et al., 2013). SRSF9 has been implicated in the control of cell invasion, not only in breast, but also in bladder-cancer cells (Yoshino et al., 2012). Finally, SRSF9-OE was previously shown to promote β-catenin accumulation and activate the Wnt signaling pathway (Fu et al., 2013), thus providing a possible link between SRSF9 expression and cell-invasion regulation. Thus, our findings reveal that specific SFs, *i.e.*, SRSF4, SRSF6, and TRA2β, play a role in both tumor initiation and metastasis, whereas others, *i.e.*, SRSF9, affect only cell invasion and metastasis. In mammary cells, then, the role of SFs in the early steps of transformation can be uncoupled from tumor dissemination and metastasis.

### The splicing repertoire of SR proteins in human mammary cells provides a functional link to their role in breast cancer

We show that mammary cells that overexpress SRSF4, SRSF6, or TRA2β exhibit similarities not only at the phenotypic level, but also at the splicing and gene-expression levels. Our analysis provides the first comprehensive repertoire of SRSF4-, SRSF6-, and TRA2β-associated DSEs in human mammary cells, and identifies unique and shared DSEs for each of SF (Figure 3 and Table S3). Significant numbers of DSEs were shared in SRSF4-, SRSF6-, and TRA2β-OE mammary cells *vs.* controls. For example, >1700 DSEs were regulated in both SRSF4-OE and SRSF6-OE cells; similarly, TRA2β-OE cells shared >1200 DSEs with SRSF4-OE cells and >1000 with SRSF6-OE cells (Figure S5 and Table S3). Moreover, this overlap increased when comparing not the specific splicing events per se, but the genes affected by alternative splicing. Oncogenic SFs that led to a similar phenotype, *i.e.*, SRSF4, SRSF6, or TRA2β, shared 16-25% of DSGs (Figure S5 and Table S3). Finally, we found that particular SFs shared substantial numbers of differentially expressed genes. SRSF4-, SRSF6- and TRA2β-OE acini exhibited a large overlap in gene-expression regulation with >250 differentially expressed genes in common (Table S3). Thus, in human mammary epithelial cells, SRSF4, SRSF6, and TRA2β regulate not only a set of shared genes, but also shared spliced isoforms that affect genes associated with biological processes involved in transformation and metastasis (Table S3).

Interestingly, several SF-induced spliced isoforms detected in this study (Figure 3 and Table S3), *i.e., ARHGEF11*+ex38, *RALGPS2*Δex10, *MLPH*+ex9, *CD44*Δex4-16, and *APLP2*Δex7, were previously associated with EMT in mammary epithelial cells (Brown et al., 2011; Harvey et al., 2018; Shapiro et al., 2011; Warzecha et al., 2010), consistent with the invasive phenotype we observe in SF-OE cells. Additionally, among the SF-induced DSEs that we validated in MCF-10A cells, several isoforms were previously linked with cell invasion or cell proliferation, *e.g.*,: i) the *ARHGEF11*+ex38 isoform was found expressed in invasive basal breast-cancer cells but not in non-invasive luminal cells, and was shown to be required for breast-cancer-cell migration and growth *in vitro* and *in vivo* (Itoh et al., 2017). The insertion of 32 amino acids coded by exon 38 abolished ARHGEF11 interaction with tight-junction protein ZO-1 and shifted ARHGEF11 localization away from cell-cell junctions (Itoh et al., 2017); ii) alternative splicing of *CD44—*and in particular a switch from the *CD44v* isoforms, which include a combination of the variable exons 6-14*—*to the *CD44s* isoforms, lacking exons 6-14, is associated with tumor progression (Chen et al., 2018). Moreover, expression of *CD44s* isoforms potentiates Akt activation and promotes cell survival in TGF-β–induced mesenchymal MCF-10A cells (Brown et al., 2011); iii) LSR expression has been implicated in breast-cancer-cell migration, and differences in its cellular localization are associated with patient outcomes (Reaves et al., 2014; Reaves et al., 2017). Skipping of exon 5, which encodes the transmembrane domain, could affect LSR localization and activity (Reaves et al., 2017); iv) *TCF7L2* encodes a protein that interacts with β-catenin and plays a role in Wnt growth-factor signaling. A polymorphism in the *TCF7L2* gene is associated with breast cancer (Chen et al., 2013b). In addition, the TCF7L2+ex4 isoform contains 23 additional amino acids and displays reduced ability to activate Wnt/β- catenin targets (Weise et al., 2010); v) The MKI67Δex7 isoform lacks 360 amino acids compared to the full-length protein, and its expression is associated with nutrient-starvation response (Chierico et al., 2017).

### TRA2β controls breast tumor initiation and metastasis

TRA2β is overexpressed in many tumor types, and a role for it in tumorigenesis has been suggested (Best et al., 2013). Our findings provide a functional characterization of the role of TRA2β in breast tumor initiation, as well as maintenance and metastasis *in vitro* and *in vivo*. TRA2β-OE promoted proliferation and increased invasiveness of non-transformed mammary epithelial cells (Figures 2 and 5). Concomitantly, TRA2β-OE led to changes in alternative splicing in target genes associated with cell division and proliferation, as well as with migration and cytoskeleton organization, cell signaling, and cholesterol metabolism (Figure 3 and Table S3). As previously described, TRA2β controls splicing of its paralog TRA2α in breast cancer cells (Best et al., 2014).

The molecular mechanisms through which TRA2β levels are altered in human breast tumors remain poorly understood. We showed that *TRA2β* is upregulated in 13% of breast tumors, but only 11% of these *TRA2β*-high tumors show amplification at the gene locus (Figure 1). Our findings suggest that *TRA2β* is a direct transcriptional target of the MYC oncogene, that both genes are frequently co-expressed in breast tumors, and functionally that their co-expression increases mammary acinar size (Figure 4). Further, we demonstrate that, 51% of *TRA2β*-high breast tumors exhibited *MYC* amplification or upregulation (Figure 4), suggesting that MYC transcriptional activation accounts for at least half of *TRA2β* alterations, similarly to *SRSF1* (Das et al., 2012). These findings, together with the results of previous studies (Das et al., 2012; David et al., 2010; Hsu et al., 2015; Koh et al., 2015), show that MYC plays a critical role in regulating the expression of several SFs linked to tumorigenesis, and provides a rationale for targeting splicing in MYC-driven tumors.

We also showed that *TRA2β-OE* tumors represent >40% of TNBCs (Figure S1), an aggressive breast cancer subtype with high metastasis incidence. We demonstrate that high levels of TRA2β are not only involved in tumor initiation, but are also required to maintain the invasive potential of metastatic human TNBC cells (Figure 5). *In vivo* TRA2β-KD significantly decreased metastatic dissemination and distant organ colonization of TNBC cells in a mouse model of breast cancer (Figure 7). Consistent with these results, TRA2β levels correlate with clinical outcomes in breast cancer patients (Figure 7). Remarkably, in 3D assays, TRA2β-KD was sufficient to reverse the invasive phenotype of TNBC cells, even after invasion had occurred (Figure 5). Thus modulation of TRA2β levels might represent a viable therapeutic option even for patients with advanced metastatic disease.

In metastatic TNBC cells, our findings revealed that TRA2β controlled splicing of target genes that play roles in cell invasion and have been associated with Wnt signaling, regulation of cell polarity, or cell-cycle and transcriptional control (Figure 6 and Table S5). For example, high TRA2β levels in both TNBC cells and human tumors were associated with splicing of *CCDC88C*, which encodes a signal transducer that regulates Wnt signaling and can trigger cancer-cell invasion and drive metastasis (Ara et al., 2016; Aznar et al., 2015; Ishida-Takagishi et al., 2012; Kobayashi et al., 2005). CCDC88C has multiple spliced isoforms that differ in their ability to enhance cell invasiveness (Dunkel et al., 2019). The isoform detected in the present study includes exon 26, which introduces a PTC, thus truncating the last 493 amino acids of the protein and producing a transcript that could potentially be degraded by NMD. Interestingly, Wnt signaling has been implicated in metastasis in TNBC cells (De et al., 2016). Additionally, we showed that TRA2β-KD altered splicing of *KMT5B*, encoding the lysine methyltransferase SUV420H1, which targets histone H4. Reduced H4K20me3 levels have been detected in human breast tumor tissues, and SUV420H1-OE was shown to suppress cancer-cell invasiveness (Yokoyama et al., 2014). Moreover, we found that TRA2β regulates splicing of exon 18 of *KIF23*, encoding a kinesin-like protein involved in microtubule formation and movement. Previously, high *KIF23* expression in tumors was shown to be associated with decreased overall survival and distant metastasis-free survival in breast-cancer patients, and KIF23-KD was shown to suppress proliferation of TNBC cells (Wolter et al., 2017). Another study showed that inclusion of exon 18 altered KIF23 cellular localization, and was associated with longer overall survival in hepatocellular carcinoma patients (Sun et al., 2015). Other TRA2β targets that we identified that are associated with cell invasion include *DOCK7*, a gene involved in cell invasion via Rac activation and implicated in the regulation of ErbB2 signaling (Yamauchi et al., 2008), and *DPHS*, which activates the RhoA signaling pathway, leading to increased cell motility and invasion *in vitro*, and increased tumor growth *in vivo* (Muramatsu et al., 2016). Additionally, several TRA2β-associated splicing targets identified in this study were previously associated with metabolism and cell signaling: *e.g.*, inclusion of *FAM126A* exon 11 leads to a C-terminally truncated protein that could potentially have distinct biological functions and affect PI4K localization (Baskin et al., 2016). PI4K is frequently amplified in breast tumors and its expression promotes multi-acinar formation in MCF-10A cells (Pinke and Lee, 2011). Moreover, we found that TRA2β also affects splicing of *GNAS* exon 3, thus regulating expression of the long and the short *GNAS* isoforms, which differ by 45 nucleotides and display differences in localization and activities (Bastepe, 2007). GNAS amplification has been associated with breast cancer, and, specifically, expression of the long isoform has been linked with enhanced cAMP signaling (Garcia-Murillas et al., 2014). The specific functions of each of the TRA2β-regulated isoforms remain to be determined. One may hypothesize that each TRA2β-regulated isoform contributes to a distinct aspect of TRA2β-mediated transformation, similarly to the targets of SRSF1 in breast cancer (Anczuków et al., 2015; Anczuków et al., 2012), and that SF-mediated transformation results from the activation of a downstream oncogenic splicing program, in the same way that an oncogenic transcription factor activates hundreds of transcriptional targets.

### RNA-binding proteins as breast-cancer drivers

This study focused on the SR protein family, as well as several related SFs. However, alterations in other RNA-binding proteins have been reported in breast tumors, and could contribute to transformation (Urbanski et al., 2018). In breast cancer, SF mutations remain rare; for example, mutations in *SF3B1* were detected in only 1.6% of TCGA breast tumors, all subtypes combined, (Seiler et al., 2018), whereas these mutations are found in 20% of uveal melanomas and 70-80% of myelodysplastic syndromes (Urbanski et al., 2018). Moreover, SF3B1 mutations are associated with luminal tumors, a breast cancer subtype that has a better prognosis than TNBC (Maguire et al., 2015). Similarly, *U2AF1, SRSF2, RBM10, QKI*, and *FUBP1* mutations are each found in less than 0.5% of TCGA breast tumors (Seiler et al., 2018).Thus, in breast cancer, SF mutations may not represent the most valuable markers for tumor progression or targets for therapeutics. On the other hand, changes in SF levels are frequently detected, and may play a role in tumorigenesis. Beyond SR proteins, others SFs are dysregulated in tumors; for example, hnRNPK, hnRNPM, hnRNPI (PTBP1), and RBM10 are upregulated in breast tumors, and both upregulated and downregulated expression of *RBM5* was reported in breast tumors (Urbanski et al., 2018). hnRNPM regulates EMT in breast epithelial cells, in part by promoting splicing of the *CD44s* isoform, and by altering TGF-β signaling (Xu et al., 2014). PTBP1-OE increases proliferation, anchorage-independent growth, and invasion in cancer cell lines (He et al., 2014; Wang et al., 2008), and PTBP1- KD in breast cancer cells had the opposite effect (Wang et al., 2018). Furthermore, PRPF8- and PRPF38A-KD, or treatment with the spliceosome inhibitor E7107, inhibits growth of TNBC cells and patient-derived TNBC xenografts (Chan et al., 2017). Whether all of these SF alterations trigger changes in a shared set of splicing targets in the mammary tissue, or affect genes converging on pathways that lead to transformation, remains to be determined. In addition, it is not yet known which of these SF alterations are drivers of tumor initiation and which are the consequence of a hyper-proliferative cell state. Our findings suggest that specific SF alterations, such as SRSF4-, SRSF6-, SRSF9-, or TRA2β-OE can contribute to tumor initiation and/or metastasis. However we also demonstrate that OE of several SFs frequently upregulated in breast tumors, such as SRSF2 or SRSF3, is not sufficient by itself to promote transformation in the model systems tested here. These findings suggest that not all SF-alterations are oncogenic in the mammary context, or that they might require additional hits to cooperate in transformation. Considering that tumors often exhibit alterations in multiple SFs, future studies should aim to understand how individual SFs regulate each other and whether they cooperate in transformation.

## AUTHOR CONTRIBUTIONS

O.A. designed and supervised the study, conducted the experiments, analyzed the data, and wrote the paper. S.D. contributed to *in vitro* and *in vivo* experiments. S.D., S.P., M.B, L.U., N.L., C.H., and M.F. contributed to cell-based assays, western blotting and RT-PCR analyses. M.A. contributed to the development of bioinformatics tools, bioinformatics analyses and bioinformatics data interpretation. A.G., A.K.K., J.G., X.H, and K.L. contributed to bioinformatics analyses. S.K.M. shared protocols and reagents. A.R.K. contributed to the study design and the paper writing. All authors discussed the results and commented on the manuscript.

## ACKNOWLEDGMENTS

We thank Stephen Samson and Taneli Helenius for helpful comments on the manuscript. We acknowledge assistance from the JAX and CSHL Microscopy and Sequencing Shared Resources, funded in part by NCI Cancer Center Support Grants P30CA034196 (JAX) and 5P30CA045508 (CSHL). This work was supported by grants from the NCI (CA1728206 to O.A and CA13106 to A.R.K.), postdoctoral awards to O.A. from the Susan B. Komen Foundation for the Cure (KG091029) and the Terri Brodeur Breast Cancer Foundation (66810101), as well as JAX start-up funds (O.A.). We acknowledge the use of data generated by TCGA, managed by NCI and NHGRI, and processed using the ISB Cancer Genome Cloud resource. The content is solely the responsibility of the authors and does not necessarily represent the official views of the National Institutes of Health.

